# Isolation and characterization of novel phages targeting pathogenic *Klebsiella pneumoniae*

**DOI:** 10.1101/2021.06.30.450654

**Authors:** Na Li, Yigang Zeng, Rong Bao, Tongyu Zhu, Demeng Tan, Bijie Hu

**Author notes:** Correspondence: Demeng Tan and Bijie Hu. These authors contributed equally: Na Li, Yigang Zeng.

## Abstract

*Klebsiella pneumoniae* is a dominant cause of community-acquired and nosocomial infections, specifically among immunocompromised individuals. The increasing occurrence of multidrug-resistant (MDR) isolates has significantly impacted the effectiveness of antimicrobial agents. As antibiotic resistance is becoming prevalent, the use of bacteriophages to treat pathogenic bacterial infections is becoming revitalized. Elucidating the details of phage-bacteria interactions will provide insights into phage biology and the better development of phage therapies. In this study, a total of 22 *K. pneumoniae* isolates were assessed for their genetic and phenotypic relatedness by multi-locus sequence typing (MLST), endonuclease S1 nuclease pulsed-field gel electrophoresis (S1-PFGE), and *in vitro* antibiotic susceptibility testing. In addition, the beta-lactamase gene (*bla*_KPC_) was characterized to determine the spread and outbreak of *K. pneumoniae* carbapenemase (KPC)-producing enterobacterial pathogens. Using these *K. pneumoniae* isolates, three phages were isolated and characterized to evaluate the application of lytic phages against these 22 ST11 *K. pneumoniae* isolates. *In vitro* inhibition assays with three phages and *K. pneumoniae* strain ZS15 demonstrated the strong lytic potential of the phages, however, followed by the rapid growth of phage-resistant mutants. Together, this data adds more comprehensive knowledge to known phage biology and further emphasizes their complexity and future challenges to overcome prior to using phages for controlling this important MDR bacterium.

## Introduction

The ongoing explosion of the antimicrobial resistance crisis has been accelerated by the overuse and misuse of antibiotics, causing the emergence and increased prevalence of MDR bacteria (Ventola, 2015). *Klebsiella pneumoniae*, a Gram-negative *Enterobacterales*, can be found in the environment and has been associated with community-acquired infections (Pendleton et al., 2013). The increasingly common mechanism of carbapenem resistance in *K. pneumoniae* is mediated by extended-spectrum β-lactamases (ESBLs), which are able to break the β-lactam structure in the most commonly used antibiotics (Pitout et al., 1997). Thus, infections of KPC-producing bacteria are consistently associated with poor clinical outcomes and represent a significant and continued threat to the healthcare system. As this problem continues to grow, there is a strong need to find an alternative means to conventional antimicrobials in order to combat this clinical pathogen.

Bacteriophages (phages for short) are the most common and diverse biological entities in the biosphere, playing an important role in promoting bacterial diversity, structuring community composition, and being a key driver in nutrient turnover (Suttle, 2005). Lytic phages infect and lyse bacterial hosts and are an attractive alternative towards combating the fast development of antibiotic-resistant *K. pneumoniae* (Sulakvelidze et al., 2001;Hung et al., 2011). The use of phages has gained interest over the past years and phages that are specific for MDR bacterial pathogens, including *Pseudomonas aeruginosa, Enterococcus* spp., *Mycobacterium abscessus, Staphylococcus aureus, A. baumannii*, and *Escherichia coli* have been identified and used to treat bacterial infections (Chibani-Chennoufi et al., 2004b;Capparelli et al., 2007;McVay et al., 2007;Letkiewicz et al., 2009;Schooley et al., 2017;Abd El-Aziz et al., 2019;Dedrick et al., 2019;Cano et al., 2020). These studies demonstrated that lytic phages can be harnessed to prevent and eradicate pathogenic bacteria, and in some cases, alleviate current medical challenges posed by antibacterial resistance and thus further reduce mortalities. While previous applications of phage therapy have shown promising results, the technology is still in a stage of development due to rapid evolution dynamics between phages and their hosts (Chibani-Chennoufi et al., 2004a). It is, therefore, crucial to characterize both phage and host communities prior to any intended phage applications.

In this study, we analyzed the molecular epidemiological feature of a collection of 22 clinical *K. pneumoniae* isolates, specifically on the characteristics of MLST, plasmid, and antibiotic susceptibility patterns. Using these *K. pneumoniae* isolates, we isolated and characterize three phages native to the hospital sewage. Details of phage host range, genome, morphological characters, and phage-host interactions were established to assess the application of phage therapies against the carbapenem-resistant isolates. Together, our study provides important information regarding the treatment and prevention of this important human pathogen.

## Materials and methods

### Bacterial strains and growth conditions

Twenty-two nonduplicate bacterial isolates of *K. pneumoniae* were collected from Zhongshan Hospital, Shanghai, China and identified with MALDI-TOF Mass Spectrometry (Bruker, USA) from June 2019 - May 2020. Isolates were routinely grown in either Luria-Bertani broth (LB) containing per liter: tryptophan 10 g, NaCl 10 g, and Yeast extract 5 g with aeration, or, on LB agar (15 g agar per liter) plates at 37°C. The beta-lactamase gene (*bla*_KPC_) was detected by PCR Master Mix (Thermo, CA, USA) with forward primer 5’-TGTAAGTTACCGCGCTGAGG-3’ and reverse primer 5’-CCAGACGACGGCATAGTCAT-3’. All strains were stored at −80°C in LB broth with 25% glycerol (vol/vol) and sub-cultivated before each use.

### Multi-locus sequence typing (MLST) and antibiotic susceptibility of bacterial isolates

In order to study population genetics and micro-evolution of this epidemic bacteria, MLST was applied to characterize these 22 isolates using sequences of internal fragments of seven house-keeping genes, according to the protocol available on the MLST Pasteur website (https://bigsdb.pasteur.fr/klebsiella/klebsiella.html) (Diancourt et al., 2005). The purified MLST amplicons were sent for Sanger sequencing, followed by further comparison with those available on the MLST database to determine the sequence type (ST). The VITEK2 compact automated system (bioMérieux, Lyon, France) was used to determine antibiotic susceptibility profiles according to the manufacturer’s instructions. The susceptibility breakpoints were interpreted with reference to the latest documents from the Clinical and Laboratory Standards Institute (CLSI).

### S1-Pulsed Field Gel Electrophoresis (S1-PFGE)

*Klebsiella* with plasmid-mediated carbapenem resistance poses significant challenges in clinical practice. To characterize the plasmid content in each strain, S1 nuclease digestion analyzed with PFGE was performed using a CHEF Mapper XA apparatus (Bio-Rad Laboratories, Hercules, CA). Briefly, the adjusted bacterial suspension mixed in a 1% agarose gel was digested with a proteinase K solution for 2 h at 54°C, then further treated with S1 nuclease. The electrophoresis conditions used in this study were as follows: initial switch time, 3 s; a final switch time, 36 s; gradient, 6 V/cm; angle, 120; temperature, 14°C; and time, 18.5 h. After the electrophoresis run was complete, the gel was stained with 1 mg/mL ethidium bromide and visualized with the ChemiDoc XRS system (Bio-Rad Laboratories, CA, USA). Genetic similarities were measured using the Dice coefficient and the clustering was carried out by the unweighted-pair group method with arithmetic averages (UPGMA) in BioNumerics (Applied Maths).

### Isolation and proliferation of phage

Phages were isolated using an enrichment protocol as described previously (Tan et al., 2014). Briefly, infiltrated hospital sewage was mixed with the same volume of 2 × LB broth and incubated at 37°C for 6 h to stimulate proliferation of total abundances of specific phages targeting the *K. pneumoniae* isolates above. Aliquots of enrichment samples were centrifuged (12,000×*g*, 10 min, 4°C) and sterilized with 0.1% chloroform. The presence of *K. pneumoniae*-phages in the hospital sewage samples was detected by double-layer agar assays with different fold dilutions of the cell-free spent supernatant from the enrichment assay. In order to obtain individual phages, single plaques were picked and purified three times with their corresponding hosts. Phage stocks were enriched and titrated using double-layer agar assays as described previously and stored at 4°C before use (Kropinski et al., 2009).

### Phage-host range assay

The host range of obtained phages was determined against 22 clinical isolates by standard spot assays. Aliquots of 2 µL diluted phage lysate (∼10^5^ PFU ml^-1^) was spotted on lawns containing 4 mL top agar mixed with 300 µL mid-log phase bacteria. After incubating at 37°C for 12 h, the spots were assessed by the turbidity of plaques and characterized as strong infection (clear), weak infection (turbid), or no inhibition (no plaque). The experiment was undertaken as independent duplicates.

### Susceptibility of strain ZS15 to phage infection and isolation of phage-resistant mutants

Overnight bacterial cultures were 1,000-fold diluted into LB broth and grown to an exponential phase with an OD600 at 0.3 and infected with its corresponding phages at an multiplicity Of infection (MOI) of 0.01. The suspension was incubated at 37°C with aeration and samples were harvested at 1-h intervals. The phage lysis efficiency was determined by optical density measurements. The experiment was done in triplicate along with parallel control cultures without phages. At time point 8-h, phage lysates were streaked onto LB agar plates and individual colonies were subsequentially purified to determine phage susceptibility by spot test assays as described above.

### Phage DNA extraction and genomic analysis

For the extraction of phage DNA, 2 mL aliquots of phage lysate with a titer of approximately 10^9^ PFU mL^-1^ was digested with DNase I (Thermo, CA, USA) and RNase A (Promega, USA) to remove any residual bacterial genomic DNA and RNA according to the manufacturer’s protocol. After digesting for 1.5 h at 37°C, EDTA (Thermo, CA, USA) was added to a final concentration of 20 mM to inactivate DNase I and RNase A. Total phage DNA was extracted using the Qiagen DNeasy Blood and Tissue kit (Qiagen, CA, USA) following the manufacturer’s instructions. Genomic DNA was sent to Sangon Biotech (Shanghai, China) and sequenced using an Illumina HiSeq sequencer.

The raw sequence containing the phage genome was quality controlled and trimmed with FastQC 0.11.2 (https://www.bioinformatics.babraham.ac.uk/projects/fastqc/) and Trimmomatic 0.36 (http://www.usadellab.org/cms/?page=trimmomatic). The trimmed reads were assembled into a single raw contig via SPAdes 3.5.0. (http://cab.spbu.ru/software/spades/). Genes were predicted and annotated using Rapid Annotation using Subsystem Technology (RAST, http://rast.nmpdr.org/). Phage virulence factor was examined using Virulence Factor Database (VFDB, http://www.mgc.ac.cn/VFs/). Phylogenetic relationship between the genome of phage NL_ZS_1, NL_ZS_2, and NL_ZS_3 and the genomes of related phages were analyzed using CLC Main Workbench (CLC, Qiagen). To construct the phylogenetic tree of phage terminase large subunits, the reference amino acid sequences of were collected from the NCBI database. The terminase large subunits were aligned, and neighbor-joining tree were created using Jukes-Cantor model with a bootstrap analysis of 1000 replicates.

### Bacteriophage morphology by TEM

Based on the genomic differences, phages were selected for examination by transmission electron microscopy (TEM). Phage stocks were back-diluted in a SM buffer (50 mM Tris-Cl, pH 7.5, 99 mM NaCl, 8 mM MgSO_4_, 0.01% gelatin) to a titer of 10^8^ PFU/mL. Diluted phage lysates were then placed on the surface of Formvar carbon-coated grids (Sigma-Aldrich, St. Louis, MO, USA) for 5 min to allow adsorption. Grids were stained with 2% sodium phosphotungstate for 2 min prior to being washed in a drop of distilled water and air-dried before being examined using a JEM-2100 microscope operated at 80 kV.

## Results

### Isolation and characterization of *K. pneumoniae* strains

The emergence of carbapenem-resistant *K. pneumoniae* (CRKP) has become one major burden of healthcare-associated infection due to the dispersion of bacteria carrying the *bla*_KPC_ gene (Bush and Bradford, 2016). KPC was first found in carbapenem-resistant *K. pneumoniae* strains in 2001 in North America, and now KPC-producing strains can be frequently isolated from nosocomial samples. It is, therefore, essential to determine and characterize the nosocomial spread of KPC clones and their corresponding phenotype and genotype. Together, 22 *K. pneumoniae* strains were recovered from various clinical specimens. According to the *K. pneumoniae* MLST database, all of the 22 clinical isolates belonged to ST11 (allelic profile 3311114), which is the predominant clone of KPC-producing *K. pneumoniae* in China (Table 1). Despite more than 100 different sequence types (STs) being reported worldwide, the mass dissemination of carbapenemase-producing *K. pneumoniae* has been restricted largely to ST258, ST11, ST340, and ST512 (Pitout et al., 2015).

**Table 1.** MLST types and antibiotic susceptibility profiles of the 22 Klebsiella pneumoniae isolates and their corresponding phage-resistant mutants. Antibiotic abbreviations: ticarcillin/clavulanic acid (TIM), piperacillin/tazobactam (TZP), ceftazidime (CAZ), cefoperazone (CPZ), cefepime (FEP), aztreonam (ATM), imipenem (IPM), meropenem (MEM), amikacin (AMK), tobramycin (TOB), ciprofloxacin (CIP), levofloxacin (LVX), doxycycline (DOX), minocycline (MIN), tigecycline (TGC), colistin (CST), trimethoprim-sulfamethoxazole (SXT).

| Strain | MLST | *bla*<sub>KPC</sub> | TIM | TZP | CAZ | CPZ | FEP | ATM | IMP | MEM | AMK | TOB | CIP | LVX | DOX | MIN | TGC | CST | SXT |
| --- | --- | --- | --- | --- | --- | --- | --- | --- | --- | --- | --- | --- | --- | --- | --- | --- | --- | --- | --- |
| ZS1 | ST11 | Yes | ≥128 | ≥128 | ≥64 | ≥64 | ≥32 | ≥64 | ≥16 | ≥16 | ≥64 | ≥16 | ≥4 | ≥8 | 2 | 4 | ≤0.5 | ≤0.5 | ≤20 |
| ZS2 | ST11 | Yes | ≥128 | ≥128 | ≥64 | 32 | ≥32 | ≥64 | ≥16 | ≥16 | 4 | 8 | ≥4 | ≥8 | 2 | 8 | 1 | ≥16 | ≤20 |
| ZS3 | ST11 | No | ≥128 | ≥128 | ≥64 | ≥64 | ≥32 | ≥64 | 1 | 1 | ≥64 | ≥16 | ≥4 | ≥8 | 4 | ≥16 | 2 | ≤0.5 | ≤20 |
| ZS4 | ST11 | Yes | ≥128 | ≥128 | 32 | ≥64 | ≥32 | ≥64 | 0.5 | ≤0.25 | ≥64 | ≥16 | ≥4 | ≥8 | 2 | 8 | 1 | ≤0.5 | ≤20 |
| ZS5 | ST11 | Yes | ≥128 | ≥128 | ≥64 | ≥64 | ≥32 | ≥64 | ≥16 | ≥16 | 4 | ≤1 | ≥4 | ≥8 | 8 | ≥16 | 4 | ≤0.5 | ≤20 |
| ZS6 | ST11 | Yes | ≥128 | ≥128 | ≥64 | ≥64 | ≥32 | ≥64 | ≥16 | ≥16 | ≥64 | ≥16 | ≥4 | ≥8 | 1 | 2 | ≤0.5 | ≤0.5 | ≤20 |
| ZS7 | ST11 | Yes | ≥128 | ≥128 | ≥64 | ≥64 | ≥32 | ≥64 | ≥16 | ≥16 | 4 | ≤1 | ≥4 | ≥8 | ≥16 | ≥16 | ≥8 | ≤0.5 | ≤20 |
| ZS8 | ST11 | Yes | ≥128 | ≥128 | ≥64 | ≥64 | ≥32 | ≥64 | ≥16 | ≥16 | ≥64 | ≥16 | ≥4 | ≥8 | 4 | 8 | 2 | ≤0.5 | ≤20 |
| ZS9 | ST11 | Yes | ≥128 | ≥128 | ≥64 | ≥64 | ≥32 | ≥64 | ≥16 | ≥16 | ≥64 | ≥16 | ≥4 | ≥8 | 4 | 8 | 2 | ≤0.5 | ≤20 |
| ZS10 | ST11 | Yes | ≥128 | ≥128 | ≥64 | ≥64 | ≥32 | ≥64 | ≥16 | ≥16 | ≥64 | ≥16 | ≥4 | ≥8 | ≥16 | ≥16 | 4 | 4 | ≥320 |
| ZS11 | ST11 | Yes | ≥128 | ≥128 | ≥64 | ≥64 | ≥32 | ≥64 | ≥16 | ≥16 | ≥64 | ≥16 | ≥4 | ≥8 | 2 | 8 | 2 | ≤0.5 | ≤20 |
| ZS12 | ST11 | Yes | ≥128 | ≥128 | ≥64 | ≥64 | ≥32 | ≥64 | ≥16 | ≥16 | ≥64 | ≥16 | ≥4 | ≥8 | 2 | 8 | 1 | ≤0.5 | ≤20 |
| ZS13 | ST11 | Yes | ≥128 | ≥128 | ≥64 | ≥64 | ≥32 | ≥64 | ≥16 | ≥16 | ≥64 | ≥16 | ≥4 | ≥8 | 2 | 8 | ≤0.5 | ≤0.5 | ≤20 |
| ZS14 | ST11 | No | ≥128 | ≥128 | 32 | ≥64 | ≥32 | ≥64 | 1 | 0.5 | ≥64 | ≥16 | ≥4 | ≥8 | 2 | 8 | ≤0.5 | ≤0.5 | ≤20 |
| ZS15 | ST11 | Yes | ≥128 | ≥128 | ≥64 | ≥64 | ≥32 | ≥64 | ≥16 | ≥16 | ≥64 | ≥16 | ≥4 | ≥8 | 2 | 8 | 8 | ≤0.5 | ≤20 |
| ZS16 | ST11 | Yes | ≥128 | ≥128 | ≥64 | ≥64 | ≥32 | ≥64 | ≥16 | ≥16 | ≥64 | ≥16 | ≥4 | ≥8 | 4 | ≥16 | 2 | ≥16 | ≤20 |
| ZS17 | ST11 | Yes | ≥128 | ≥128 | ≥64 | ≥64 | ≥32 | ≥64 | ≥16 | ≥16 | ≥64 | ≥16 | ≥4 | ≥8 | 4 | 8 | 1 | ≤0.5 | ≤20 |
| ZS18 | ST11 | Yes | ≥128 | ≥128 | ≥64 | 32 | ≥32 | ≥64 | ≥16 | 8 | 8 | ≤1 | ≥4 | ≥8 | ≤0.5 | 2 | ≤0.5 | ≤0.5 | ≤20 |
| ZS19 | ST11 | No | ≥128 | ≥128 | 8 | ≥64 | ≥32 | ≥64 | ≤0.25 | ≤0.25 | ≥64 | ≥16 | ≥4 | ≥8 | 2 | 4 | ≤0.5 | ≤0.5 | ≤20 |
| ZS20 | ST11 | Yes | ≥128 | ≥128 | ≥64 | ≥64 | ≥32 | ≥64 | ≥16 | ≥16 | ≥64 | ≥16 | ≥4 | ≥8 | 1 | 4 | ≤0.5 | ≤0.5 | ≤20 |
| ZS21 | ST11 | Yes | ≥128 | ≥128 | ≥64 | ≥64 | ≥32 | ≥64 | ≥16 | ≥16 | ≥64 | ≥16 | ≥4 | ≥8 | 1 | 4 | ≤0.5 | ≤0.5 | ≤20 |
| ZS22 | ST11 | Yes | ≥128 | ≥128 | ≥64 | ≥64 | ≥32 | ≥64 | ≥16 | ≥16 | ≥64 | ≥16 | ≥4 | ≥8 | 1 | 2 | ≤0.5 | ≤0.5 | ≤20 |

In addition, the *bla*_KPC_ gene was detected in 19 out of 22 isolates, indicating the prevalence of β-lactamase nosocomial pathogens (Table 1). The *bla*_KPC_ gene can be disseminated between species by horizontal (lateral) gene transfer (HGT) via various means, such as plasmids, transposons, prophages, and integrative conjugative elements (ICEs), which play an important effect in generating genotypic heterogeneity and phenotypic variation among bacterial populations (Hardiman et al., 2016;Botelho and Schulenburg, 2020). For instance, transfer of interspecies KPC can occur through the dissemination of mobile genetic elements between strains of *S. aureus* (Goren et al., 2010).

Since pathogenicity and antimicrobial resistance genes are mainly coded by plasmids, in order to identify the subtypes of *K. pneumoniae* ST11, pulsed-field gel electrophoresis with S1 nuclease (S1-PFGE) were utilized to resolve the prevalence of plasmid fingerprints of all the 22 isolates, particularly large ones, which clustered into 6 primary clades (A, B, C, D, E, and F) (Figure 1). Clade A and B were the most common types, while the other clades (C, D, E, and F) accounted for 31.8%. All groups harbored one to two plasmids ranging from 20 Kbp and 250 Kbp. Thus, it appears that S1-PFGE is a highly discriminative molecular typing method capable of effectively discriminating ST11 clones into several different subtypes, indicating the prevalence of ST11 was not clonally homogenous.

**Fig. 1.**
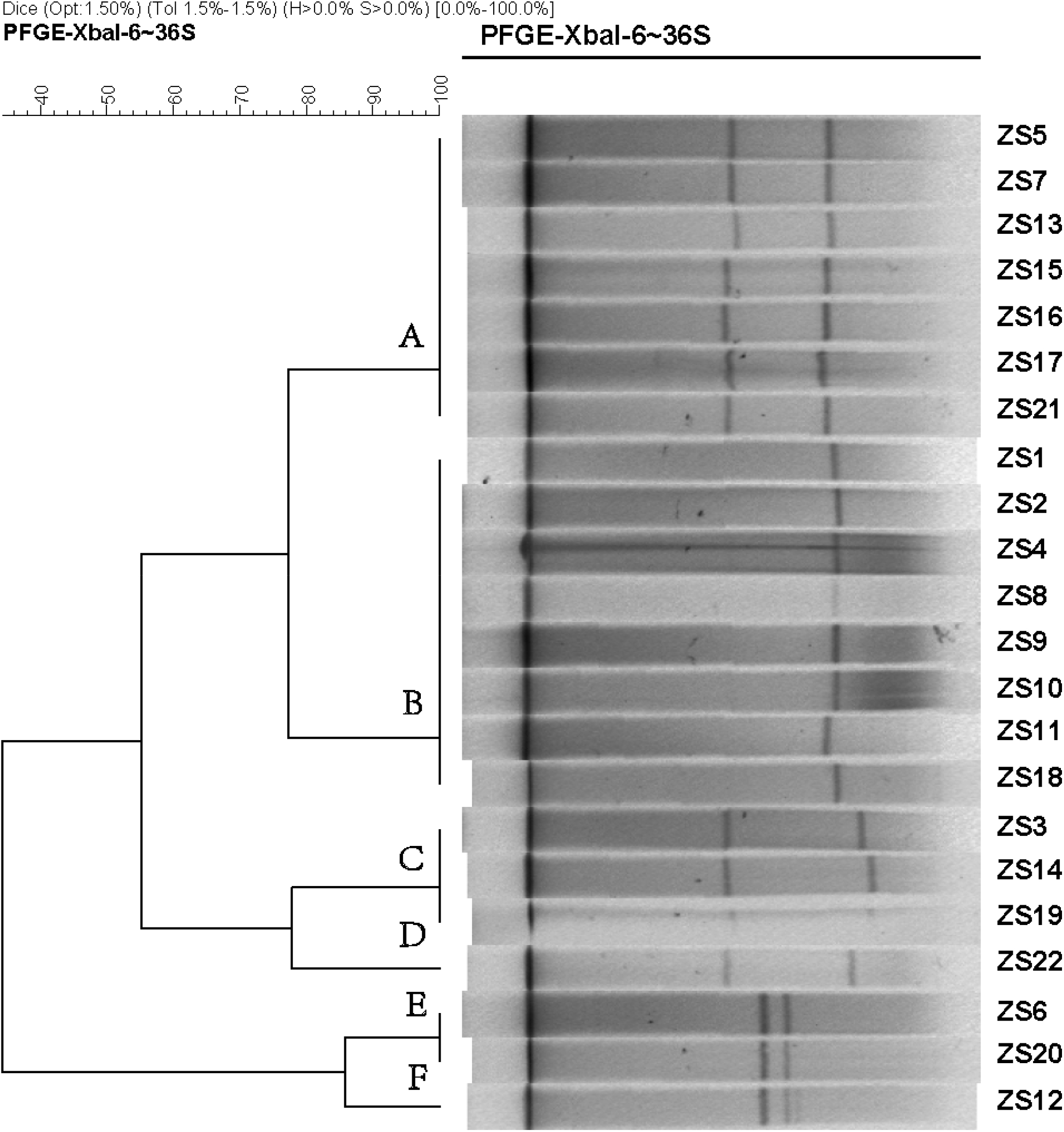
Dendrogram generated by BioNumerics software (Version7.1, Applied Maths, St-Martens-Latem, Belgium) showing cluster results of molecular epidemiology S1-PFGE analysis of 21 *K. pneumoniae* isolates. Band-based similarity coefficients were calculated using Dice coefficient and unweighted pair group average method algorithm (UPGMA).

### Imipenem and meropenem resistance involve the production of β-lactamase

Individual antimicrobial data can be found in Table 1. For ticarcillin/clavulanic acid, piperacillin/tazobactam, cefepime, aztreonam, ciprofloxacin, and levofloxacin, drug resistance was detected in all strains. In addition, low percentages of resistance to doxycycline, minocycline, tigecycline, colistin, and trimethoprim/sulfamethoxazole were also observed for most strains. Of these, 17 out of the 22 isolates become resistant to both imipenem and meropenem with a MIC of ≥16 µg/mL, while the other 5 were both sensitive to imipenem and meropenem, except for strain ZS18 which was resistant to imipenem but intermediate to meropenem. For these 5 sensitive isolates, 2 of them were confirmed by PCR and sequencing to have the *bla*_KPC_ gene. Interestingly, strain ZS4 harboring *bla*_KPC_ was susceptible to imipenem and meropenem, perhaps indicating the resistance is associated with changes in gene expression at the mRNA levels (Senchyna et al., 2019).

### Isolation and characterization of phages

Three phages (NL_ZS_1, NL_ZS_2, and NL_ZS_3) were isolated from local hospital sewage using the enrichment protocol to evaluate the potential of phage therapy as an effective alternative against *K. pneumoniae* strains. The host range of the three phages and their lytic efficiency against all of the 22 bacterial isolates were assessed using the spot test assay. According to the host range profile, the phages were able to infect all of the 22 bacterial isolates, however, the phages displayed a unique profile in terms of host range patterns and lytic efficiency (Figure 2). Based on their host susceptibilities, phages were divided into two major groups. Specifically, phage NL_ZS_3 displayed the broadest host range, while phage NL_ZS_1 and phage NL_ZS_2 showed a relatively narrow host range, infecting only ∼50% of the tested *K. pneumoniae* strains (Figure 2). The host range differences between the phages observed in this study are very likely due to the different phage characterization, i.e., efficiency of adsorption and lytic efficiency in its potential hosts.

**Fig. 2.**
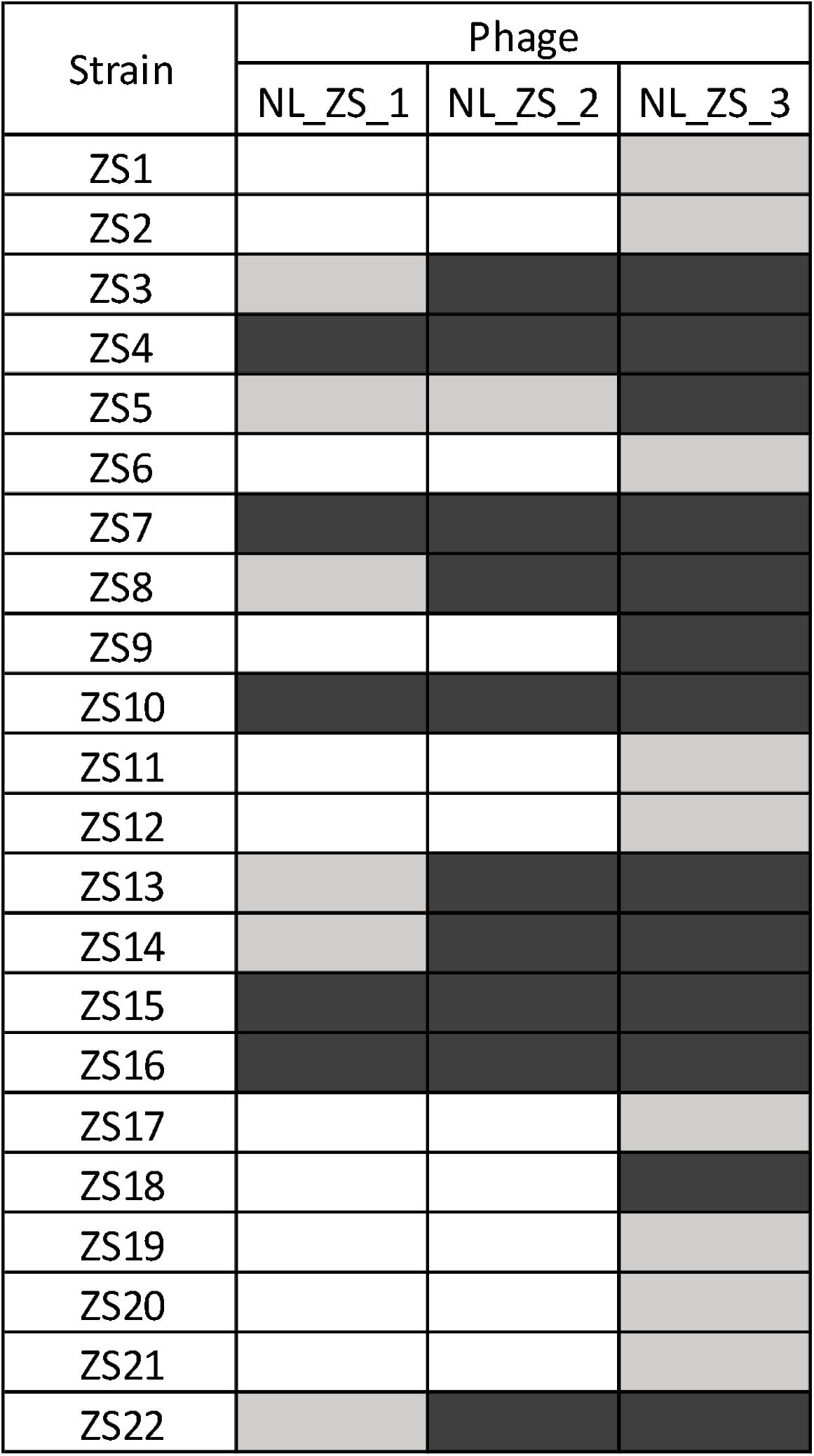
Phage-host range assay of the 3 phages (columns) against the 22 *K. pneumoniae* isolates (rows). *Klebsiella pneumoniae* phages displayed broad and narrow host ranges. Black, grey and white indicate infection, weak infection and no plaque formation, respectively.

### Morphology and genome sequence of phages

All phage lysates formed transparent plaques on strain ZS15, which are considered as an indicator of strong lysis. Plaque morphologies of all phages presented a zone of clear lysis on the host strain ZS15, while a halo zone was detected on the plaques of both phages NL_ZS_1 and NL_ZS_2, indicating the production of polysaccharide depolymerase (Figure 3). Depolymerase activity are usually encoded by their tail fibers or tail spikes protein on the baseplate, which has the potential to provide an alternative treatment to conventional antibiotics.

**Fig. 3.**
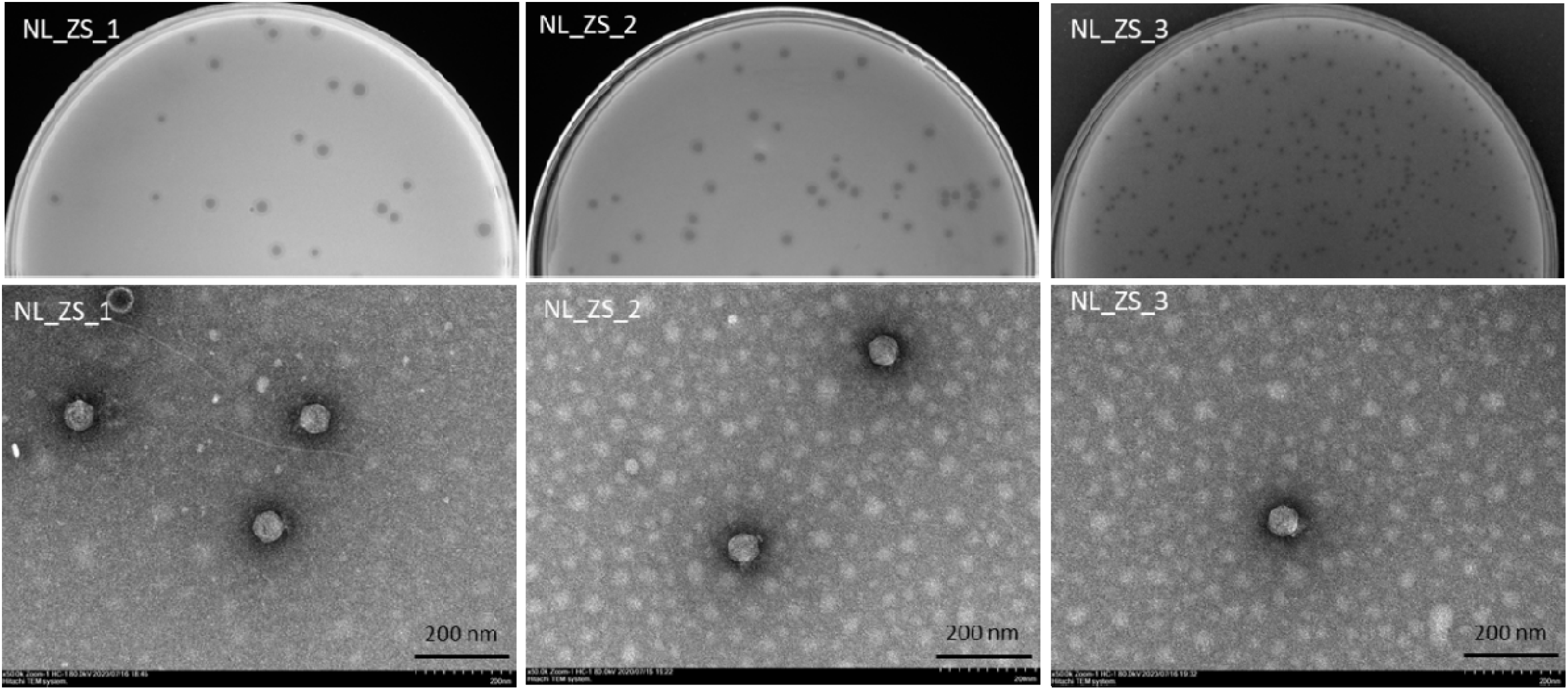
Plaque images (up) and their corresponding TEM micrographs (below). Phages were stained with 2% sodium phosphotungstate and were found to have short small podophage morphology. Scale bar, 200 nm.

Based on transmission electron microscopy (TEM) analysis, all the phages belong to the family *Podoviridae* of *Caudovirales* order based on the International Committee on Taxonomy of Viruses (ICTV) definitions, having short, non-contractile tails (∼11 nm in length) and icosahedral heads (∼61 nm diameter) (Figure 3 and Table 2). In addition, total genomic DNA was extracted from each of the three phages, and their genomes were sequenced by Illumina MiSeq. The assembly sizes of these three phages ranged from 38 kb to 40 kb, having a G+C content ranging between 50% to 53%, with 45 to 49 predicted protein-coding genes (Table 2). All phage genomes (GenBank accession: phage NL_ZS_1, MT813140; phage NL_ZS_2, MT813141; and phage NL_ZS_3, MT813142) were submitted to RAST (Rapid Annotation using Subsystem Technology) and further annotated by the available pipeline. No identifiable virulent genes, antibiotic resistance genes, or genes involved in lysogenic cycles were detected. Therefore, it appears that those phages isolated in this study can satisfy the safety assessment based on their complete genome sequences. Progressive multiple genome alignments were analyzed using Easyfig software (https://mjsull.github.io/Easyfig/) (Figure 4) to determine the relatedness of three *K. pneumoniae* phages. A considerable relation was shown between phage NL_ZS_1 and phage NL_ZS_2, while phage NL_ZS_2 displays a relatively low homologous region with phage NL_ZS_3. Likely, phylogenetic analysis of the large subunit of the terminase of phage NL_ZS_1, NL_ZS_2, and NL_ZS_3 reveals their close relation to large terminase subunits of other phages, specifically, phage NL_ZS_1 and phage NL_ZS_2 display high similarities to *Klebsiella* phage K11, BIS33, and KpV289, while phage NL_ZS_3 was closely related to *Salmonella* phage phiSG-JL2 (Figure 5). Together, these results showed high relation to the phage-host range pattern, demonstrating phage susceptibility correlates with phage core genome, such as genes involved in the phage tail fibers. However, further studies are needed to probe the genetic determinants that contribute to the host-range difference among these three phage isolates.

**Table 2.**
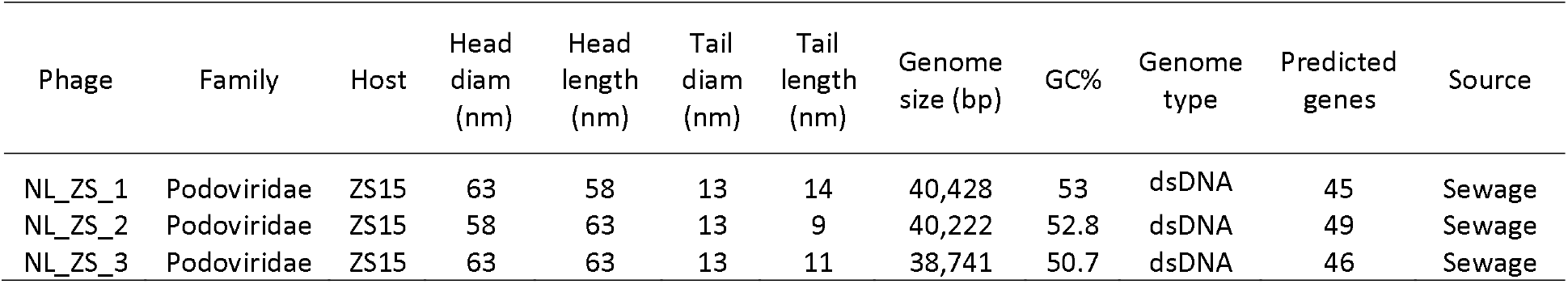
Morphological and genomic characterization of phages isolated from the local hospital sewage.

**Fig. 4.**
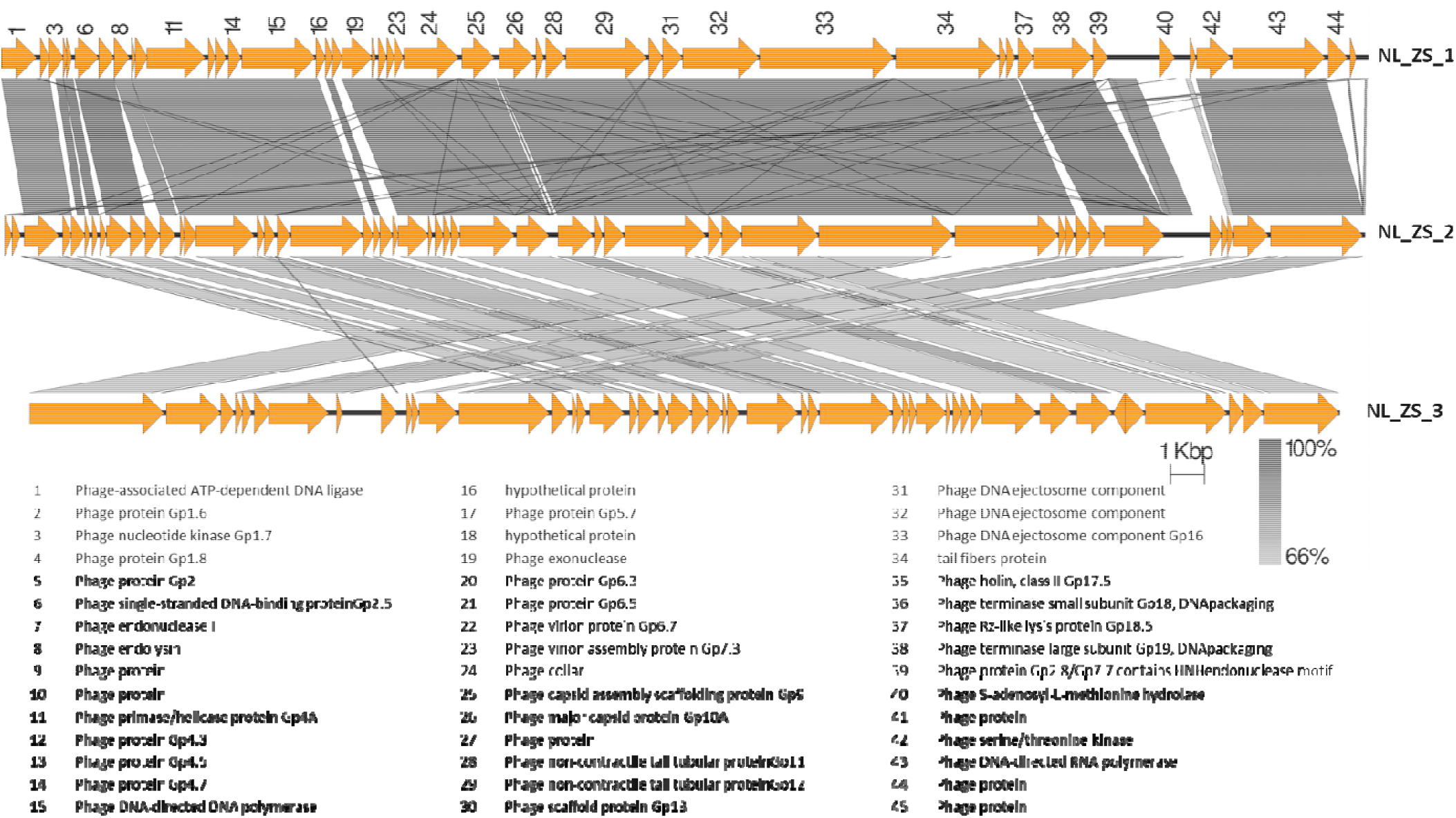
The comparison of the whole genome sequences among three *K. pneumoniae* phages (NL_ZS_1, NL_ZS_2, and NL_ZS_3) with Easyfig using the BLASTN algorithm. Grey lines connect regions indicates nucleotide identity >66%, and are gradient-colored according to their similarity; darker shade of grey represents higher identify. The designations of the open reading frames of NL_ZS_1 are shown on the top.

**Fig. 5.**
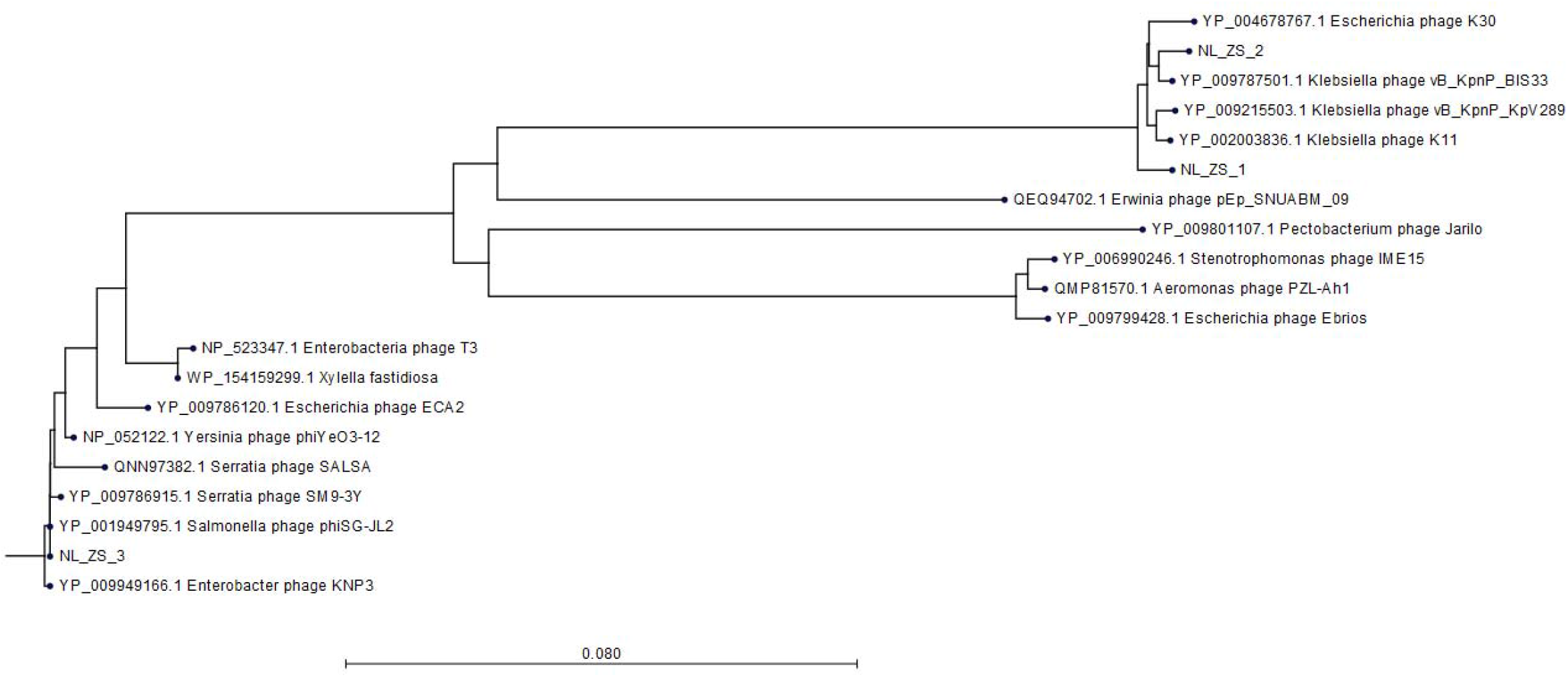
Phylogenetic tree was constructed based on the amino acid sequence of the larger terminase subunit of phages NL_ZS_1, NL_ZS_2, and NL_ZS_3 and phages sharing homology sequence identity retrieved from GenBank (https://www.ncbi.nlm.nih.gov/). The evolutionary history of 20 larger terminase subunit amino acid sequences were aligned and inferred using Neighbor Joining method and 1000 bootstrap replicates (CLC Genomics Workbench 8).

### Phage growth characteristics in LB broth

To examine the efficiency of phage infection, *K. pneumoniae* ZS15 was infection in liquid cultures with individual phages (NL_ZS_1, NL_ZS_2, and NL_ZS_3) at an MOI of ∼0.01, respectively, and their inhibition was assessed by bacterial growth following the measurement of the optical density of the bacterial cultures with and without phages. A similar inhibitory effect of phage NL_ZS_1 and phage NL_ZS_2 was observed in the *in vitro* inhibition assay with a drastic decline in optical density on the initial control of the host population for 1.5 h followed by regrowth of the resistant population (Figure 6). A similar trend in the phage-amended cultures of ZS15+ NL_ZS_3 was also observed in the first 5 h, however, it was followed by less pronounced decrease and finally stabilized after 6 h (Figure 6), After 8 h, the OD600 nm of the phage-amended cultures of group 1 (ZS15+phage NL_ZS_1 and ZS15+phage NL_ZS_2) and group 2 (ZS15+phage NL_ZS_3) stabilized at 1.96 and 0.09, corresponding to 52% to 96% of the final OD in the control group, respectively (Figure 6). Together, these results demonstrate that phage NL_ZS_3 was slightly more effective in reducing bacterial densities than the rest of the phages.

**Fig. 6.**
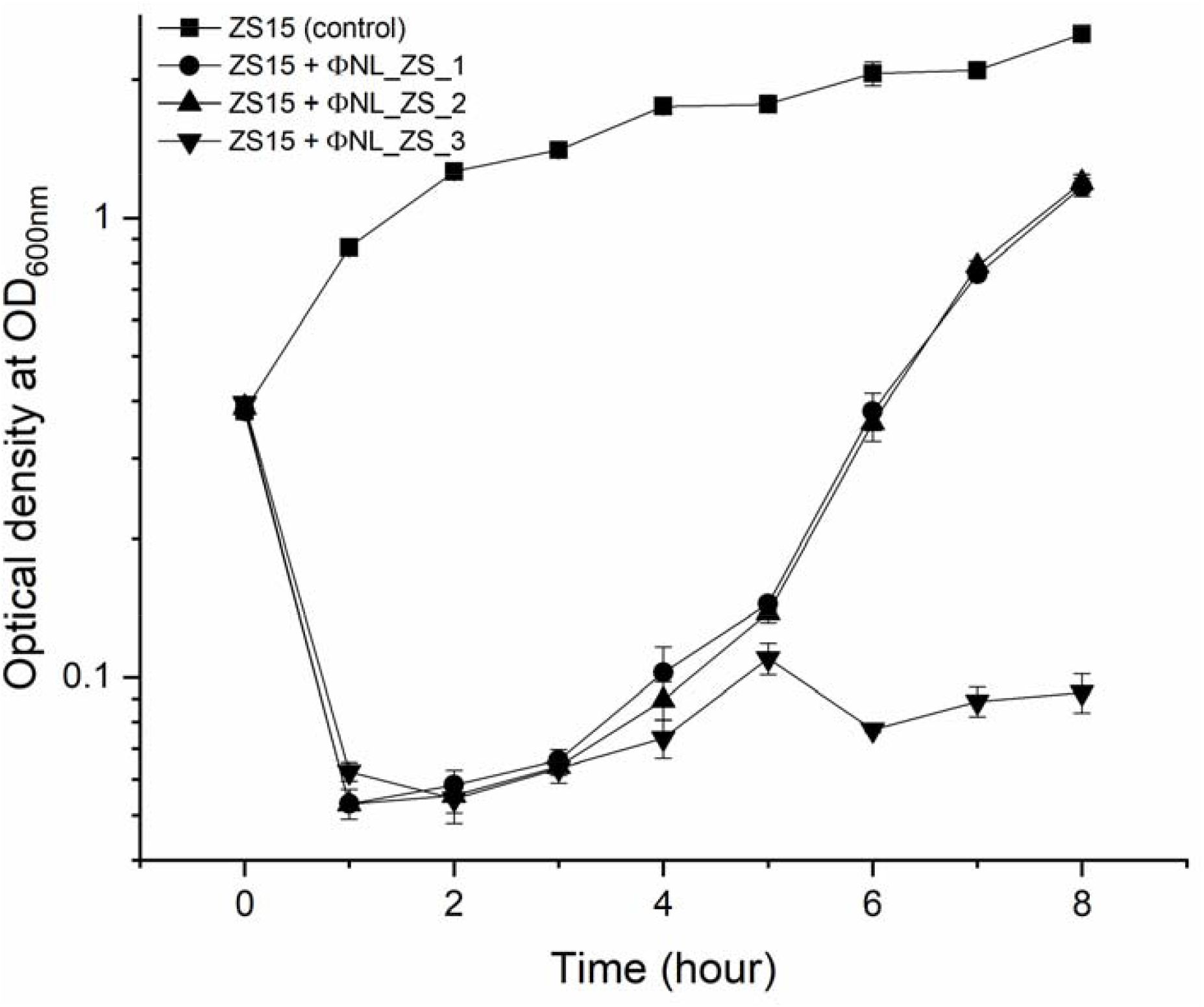
Inhibition assay of phage NL_ZS_1, NL_ZS_2, and NL_ZS_3 in the treatment at an MOI of 0.1 and control without phage in cultures of *K. pneumoniae* ZS15. Optical density (OD600) was measured at 1-h intervals over an 8-h incubation period. Error bars represent the standard deviations from all experiments, carried out in triplicate.

Despite *K. pneumoniae* ZS15 displaying a strong susceptibility to infection by all three phages, phage NL_ZS_3 was better at suppressing resistance to host strain ZS15 over phage NL_ZS_1 and NL_ZS_2. Bacteria are known to rapidly develop phage resistance when subjected to predation by lytic phages in a laboratory medium (Labrie et al., 2010). By screening phage resistance mutants, we attempted to better understand the inhibition differences among each of the phage-amended cultures. Phage insensitive mutants were isolated from phage lysate cultures during the incubation for the analysis of changes in their susceptibility upon phage selection. Based on the results of spot test assays, we observed that phage-amended cultures (ZS15+phage NL_ZS_1, ZS15+phage NL_ZS_2, and ZS15+phage NL_ZS_3) were dominated by resistant non-mucoid clones after 8 h of incubation, indicating conversion of mucoid *K. pneumoniae* strain ZS15 to non-mucoid variants by consequence of selective pressure operated by phages (Table 3). However, in the ZS15+phage NL_ZS_3 cultures, heterogeneity in phage susceptibility was observed (Table 3). Out of 30 *K. pneumoniae* ZS15 isolates, 2 (∼6.7%) were fully sensitive to phage NL_ZS_3 and displayed the hypermucoviscosity (HV) phenotype as the parental strain ZS15, and partially sensitive (i.e., reduced phage susceptibly when compared to the parental strain) subpopulations (∼50%) were isolated as well and did not revert back to the wild-type phenotype despite three rounds of purification. Interestingly, 43.3% were fully resistant to phage NL_ZS_3, however, a small fraction (∼30%) of phage NL_ZS_3 resistant mutants reverted back to the parental phenotype after three rounds of purification, suggesting different anti-phage mechanisms were allowing coexistence of phage sensitive mutants and that their lytic phages could exist between bacterial host ZS15 and phage NL_ZS_3. Since phage susceptibility was only measured at 8 h, we do not know the dynamics of phenotype heterogeneity over time or how long such persistence can last before heritage mutations dominate the entire population.

**Table 3.**
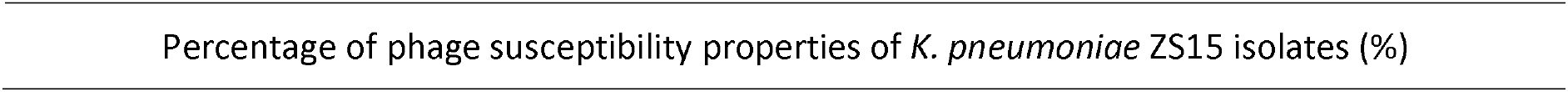
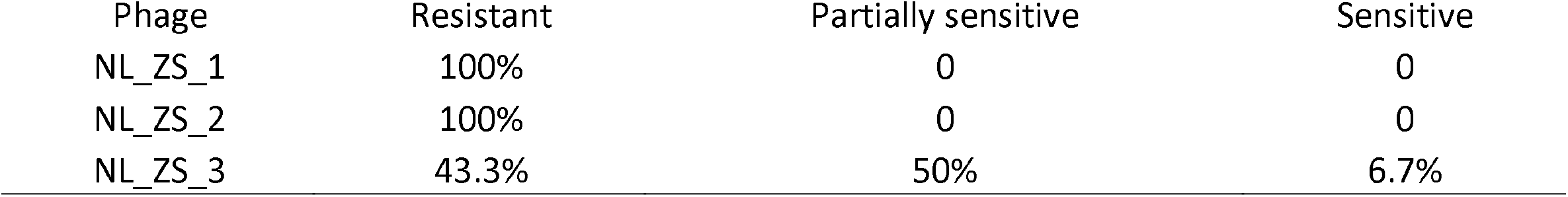
Phage susceptibility properties of K. pneumoniae ZS15 isolates obtained from cultures enriched with the phages. “Resistant” indicates that isolates were not susceptible to phage infection in the spot test. Partially sensitive” indicates that isolates had a reduced susceptibility to the phage. Sensitive” indicates that isolates were fully sensitive to the phage.

Percentage of phage susceptibility properties of *K. pneumoniae* ZS15 isolates (%)
| Phage | Resistant | Partially sensitive | Sensitive |
| --- | --- | --- | --- |
| NL_ZS_1 | 100% | 0 | 0 |
| NL_ZS_2 | 100% | 0 | 0 |
| NL_ZS_3 | 43.3% | 50% | 6.7% |

### Anti-phage defense-mediated altered sensitivity to antibiotics

Previous studies have shown that evolutionary phage-host interactions affect phage resistance and antibiotic susceptibility both positively and negatively, and are highly genotype-dependent in bacterial populations (Comeau et al., 2007;Chan et al., 2016;Burmeister et al., 2020). We started to question whether phage-resistant mutants changed their antibiotic susceptibilities via antagonistic coevolution between hosts and parasites in LB broth. To investigate this further, we performed Vitek 2 assays for antimicrobial susceptibility, testing phage-resistant mutants (ZS15-3R) and their parental phage susceptible strains (ZS15). Interestingly, for tigecycline susceptibility testing of carbapenem-resistant *K. pneumoniae* ZS15 strain, we noticed that strain ZS15 evolved resistance to phage NL_ZS_3, while simultaneously becoming more sensitive to tigecycline with the minimum inhibitory concentration shifting from 8 µg/mL to 2 µg/mL. Unfortunately, we were unable to identify potential genes related to phage selection and antibiotic sensitivity during the interactions between strain ZS15 and phage NL_ZS_3. Further, comparative genomic sequence analyses are needed to elucidate the underlying mechanisms.

## Discussion

*Klebsiella pneumoniae* is a common nosocomial pathogen that causes difficult-to-treat infections worldwide. The prevalence and distribution of *bla*_*KPC*_ gene in *K. pneumoniae* is increasing, making carbapenem resistance isolates difficult to treat due to the lack of effective and safe alternative options. Successful application of phage therapy in treating *K. pneumoniae* caused infections requires a detailed understanding of the phenotypic and genotypic diversity printers of *K. pneumoniae* pathogens as well as well-characterized phages that covers host diversities. In this study, we characterized 3 phages and 22 *K. pneumoniae* strains, covering considerable temporal variability. Further, *K. pneumoniae* strain ZS15 was selected for more detailed studies of phage lytic potential and phage-host interactions.

### Diversity of *K. pneumoniae* ST11 isolates

The clonal relationship between *K. pneumoniae* strains examined in the present study belonged to ST11 irrespective of time or place of isolation. ST11 is among the most important clinical dominant in Asia and South America and has been reported in different regions worldwide since its emergence (Qi et al., 2011). In 2018, a fatal outbreak of ST11 carbapenem-resistant hypervirulent *K. pneumoniae* in a Chinese hospital has caused high mortality and morbidity (Gu et al., 2018;Yao et al., 2018). Likely, ST258, single-locus variants of ST11 (the *tonB* allele distinguishes the two sequence types), has emerged as an important cause of hospital death in the United States and other countries, and is largely responsible for the global spread of KPC (Toth et al., 2010). However, the molecular linage between ST11 and ST258 largely remains unknown, recent comparative genomic analyses showed ST258 is a hybrid with ∼80% of the chromosome homology to ST11 strains, while the remaining 20% display high similarity to the ST442 isolates (Chen et al., 2014). Despite MLST does not provide sufficient discrimination for the 22 pathogenic isolates or resolve differences across spatial and temporal sac ales, understanding the molecular evolutionary history of bacterial isolates is an important step towards the development of diagnostic and potential therapeutic strategies to combat infections caused by multidrug-resistant *K. pneumoniae*.

Carbapenem resistance was seen among all *Klebsiella* isolates. Since carbapenem-resistant *K. pneumoniae* (CRKp) are endemic to local hospitals, where colistin has been largely used as the last treatment option to fight against these infections. Out of 22 strains, 2 isolates (strain ZS2 and ZS16) display resistance to colistin. Together, there 22 isolates could be regarded as a real superbug that could pose a major public health challenge. There is currently little understanding of molecular mechanisms underlying antibiotic resistance and its impact on virulence remain unclear, despite ∼86% of strains were confirmed to be carbapenemase gene positive. A recent sequence of complete *K. pneumoniae* genomes showed that genes coding for resistance to antibiotics or virulence factors is located in the plasmid. Moreover, the spread of these resistant plasmids into the normal gut flora poses an even great public health threat. In order to recapitulate the contents and diversity of the plasmids of genetically related *K. pneumoniae* strains, it is important to track the spread of its plasmids between each isolate. In our study, S1-PFGE showed four clusters of the CRKp producing strains that corresponded to ST11, which demonstrates a high degree of genetic diversity. Future work will seek to understand the dissemination of these plasmids and their potentials of converting any *K. pneumoniae* strains with no existing antimicrobial resistance or virulence determinants into CRKp strain immediately. Together, these results provide important insights into relatively narrow phylogenetic and phenotypic groups of bacteria, emphasizing the complexity for the development of alternatives to treat this pathogen.

### Phage isolation and diversity

Generally, phages are widespread and genetically diverse in almost all environments, especially in sewage, known as a fertility source for many pathogens. The high occurrence of *K. pneumoniae* phages in the collected samples (∼72%) suggested the *K. pneumoniae* phages are widespread in the local hospital sewage supply. The phages isolated from the local hospital sewage displayed two distinguished host ranges when tested against the collection of 22 *K. pneumoniae* isolates. Only 1 morphological family, *Podoviridae*, was represented, and the phages ranged in genome size from 38 kb to 40 kb. Due to the limited sample numbers, we do not know to what extent these isolated phages represent the true diversity and structure of local hospital sewage phage communities. Further studies of viral metagenomics that are not dependent on cell culture approaches are needed to expand our understanding of phage genomic diversity from the evolutionary perspective and perhaps genetic exchanges in response to the selection pressure from the bacterial host.

Phage NL_ZS_3 displayed a relatively broad host range, while phage NL_ZS_1 and phage NL_ZS_2 shared identical relatively narrow host ranges, suggesting that phage NL_ZS_1 and phage NL_ZS_2 might use similar phage binding receptors to recognize bacterial phage receptors in their susceptible hosts. This is interesting as bioinformatic analysis showed that phage-binding receptors of phage NL_ZS_1 and phage NL_ZS_2 were identical based on the amino acid sequence. The possible explanation for host range difference could be phage NL_ZS_3 used a conservative apparatus as its phage receptors, such as outer membrane proteins, lipopolysaccharides, flagella, or pili. Moreover, several copies of genes encoding phage tails or tail fibers were present in the phage NL_ZS_3, which opens the possibility that phage NL_ZS_3 may use a primary receptor to initiate attachment but require a secondary cell surface receptor to irreversibly bind to the bacterial host for injection, permitting more flexibility to develop on a wider range of hosts. For example, the T4 phage uses its long tail fibers to recognize lipopolysaccharide and the outer membrane protein C, present on the bacterial surface (Tétart et al., 1996;Baslé et al., 2006). This reversible attachment allows the phage to move along the surface and find a suitable site to inject its DNA.

### Phage-host interactions in strain ZS15

Isolation and characterization of bacterial isolates during the LB broth experiments confirmed the rapid development of phage resistance and subsequently dominated the bacterial populations in the phage NL_ZS_1 and phage NL_ZS_2, which is regarded as a significant barrier to the successful application of phage therapies (Koskella and Meaden, 2013). Interestingly, in the current experiment with strain ZS15, phage NL_ZS_ 3 exposures led to the rapid development of phenotypic diversity (i.e., resistant, partially sensitive, and resistant) within a population that enables survival of fluctuating, suggesting different anti-phage mechanisms were allowing coexistence of phage sensitive mutants and that their lytic phages could exist. Since phage susceptibility was only measured at 8 h, we do not know the dynamics of phenotype heterogeneity over time or how long such persistence can last before heritage mutations dominate the entire population. It is, therefore, crucial to characterize the phage-host interaction regarding the phenotypic and genetic diversification in bacteria and phages prior to any intended phage applications. The exact resistant mechanism by which strain ZS15 avoids phages infection, however, remains to be discovered.

A common form of developing phage resistance for bacterial hosts is associated with loss or altered structure of the receptors to block the potential phage adsorption in order to avoid infection (Labrie et al., 2010). Such changes are directly linked to the key step in recognition between phage receptor-binding proteins (RBP) located on tail fibers, and phage receptors encoded on the susceptible bacterial host cell surface (Bertozzi Silva et al., 2016). In general, the fitness costs resulting from receptor mutation (i.e., motility, nutrient transport, or antimicrobial susceptibility) make the mutants unable to compete with phage-susceptible strains in both environmental samples and infection models, even in the presence of phage predation (Seed et al., 2012;Sumrall et al., 2019). Understanding the evolutionary implication of phage-receptor interactions will be beneficial for predicting how phages interact with their host, as phage receptor mutation-mediated phenotypic variabilities may result in exopolysaccharide associated fitness cost. Such characterization of phage resistance evolution will be used as a guide for selecting proper phage targeting of specific receptors with a significant clinical application. Our data reveal when phage resistance evolves, the resistant mutants showed increased tigecycline sensitivity, despite the mechanism underlying potential is not fully understood in this study.

Tigecycline, known as a tetracycline antibiotic, has shown the greatest antimicrobial activity against CRKP. Genes e.g. *tet*(B), *tet*(A), *tet*(K), *tet*(M) and *tet*(S) coding for multidrug efflux pumps are membrane-related proteins and can efflux tetracyclines from the bacteria intracellularly, thus decreasing the MIC and subsequently allowing protein synthesis without affecting cell viability (Linkevicius et al., 2016). Using membrane-related proteins as a surface structure is one of the most common components functioning as a phage receptor. Therefore, it makes sense that phage-resistant defense strategies would confer a fitness cost for bacteria, i.e., becoming more sensitive to certain antibiotics. For instance, in *P. aeruginosa*, phage OMKO1 evolved to become less resistant to antibiotics. Likewise, in *E. coli*, the TolC protein evolved in a variety of diverse cellular functions, including antibiotic efflux and phage TLS receptor (Chan et al., 2016). However, unlike previous studies, the anti-phage TLS mediated TolC mutation did not influence its role in antibiotic efflux. Overall, these results reveal that a phage-host arm race can result in different outcomes in terms of antibiotic susceptibilities, highlighting the diversity and complexity of host interactions within bacterial populations.

### Potential applications of phage therapy

Successful application of phage therapy requires collaboration between microbiology and preclinical development; however, little is known about their alignment to guide future development. Since the discovery of phages, many phages have been isolated and identified as potential alternatives to antibiotics with different efficacy profiles. Phage specificity is both a benefit and an obstacle for clinical trials, as phage receptor diversity affects phage-host interaction. When the bacterial population is under phage attack, only a small fraction of cells can survive these selective pressures, and those that survive determine the fate of the population. Therefore, elucidating details of phage-host interactions and how these would influence bacterial physiological heterogeneity is fundamental to the development of phage therapies. In this study, the three highlighted phages showed promising *in vitro* effects that perhaps these phages can be tamed for our benefits without negatively affecting their natural balance. The main limitation of this study is that only a small number of bacterial isolates and phages have been investigated. A large-scale study on the interactions between carbapenem-resistant K pneumoniae strains and phages covering a considerable spatial and temporal variability are essential for the successful application of phage-based pathogen control to fill our knowledge gap between *in vitro* and *in vivo* models.

## Acknowledgments

We thank Xiao Wang and Ruoyu Li for technical assistance. We thank all members from Zhu’s lab for thoughtful discussions and grateful to Brett C. Gonzalez for proofreading the manuscript.

## References

Abd El-Aziz, A.M., Elgaml, A., and Ali, Y.M. (2019). Bacteriophage Therapy Increases Complement-Mediated Lysis of Bacteria and Enhances Bacterial Clearance After Acute Lung Infection With Multidrug-Resistant Pseudomonas aeruginosa. J Infect Dis 219, 1439–1447.

Baslé, A., Rummel, G., Storici, P., Rosenbusch, J.P., and Schirmer, T.J.J.O.M.B. (2006). Crystal structure of osmoporin OmpC from E. coli at 2.0 Å. 362, 933–942.

Bertozzi Silva, J., Storms, Z., and Sauvageau, D. (2016). Host receptors for bacteriophage adsorption. FEMS Microbiol Lett 363.

Botelho, J., and Schulenburg, H. (2020). The Role of Integrative and Conjugative Elements in Antibiotic Resistance Evolution. Trends Microbiol.

Burmeister, A.R., Fortier, A., Roush, C., Lessing, A.J., Bender, R.G., Barahman, R., Grant, R., Chan, B.K., and Turner, P.E.J.P.O.T.N.a.O.S. (2020). Pleiotropy complicates a trade-off between phage resistance and antibiotic resistance. 117, 11207–11216.

Bush, K., and Bradford, P.a.J.C.S.H.P.I.M. (2016). β-Lactams and β-lactamase inhibitors: an overview. 6, a025247.

Cano, E.J., Caflisch, K.M., Bollyky, P.L., Van Belleghem, J.D., Patel, R., Fackler, J., Brownstein, M.J., Horne, B.A., Biswas, B., and Henry, M.J.C.I.D. (2020). Phage Therapy for Limb-threatening Prosthetic Knee Klebsiella pneumoniae Infection: Case Report and In Vitro Characterization of Anti-biofilm Activity.

Capparelli, R., Parlato, M., Borriello, G., Salvatore, P., and Iannelli, D. (2007). Experimental phage therapy against Staphylococcus aureus in mice. Antimicrob Agents Chemother 51, 2765–2773.

Chan, B.K., Sistrom, M., Wertz, J.E., Kortright, K.E., Narayan, D., and Turner, P.E. (2016). Phage selection restores antibiotic sensitivity in MDR Pseudomonas aeruginosa. Sci Rep 6, 26717.

Chen, L., Mathema, B., Pitout, J.D., Deleo, F.R., and Kreiswirth, B.N.J.M. (2014). Epidemic Klebsiella pneumoniae ST258 is a hybrid strain. 5, e01355–01314.

Chibani-Chennoufi, S., Bruttin, A., Dillmann, M.L., and Brussow, H. (2004a). Phage-host interaction: an ecological perspective. J Bacteriol 186, 3677–3686.

Chibani-Chennoufi, S., Sidoti, J., Bruttin, A., Kutter, E., Sarker, S., and Brussow, H. (2004b). In vitro and in vivo bacteriolytic activities of Escherichia coli phages: implications for phage therapy. Antimicrob Agents Chemother 48, 2558–2569.

Comeau, A.M., Tétart, F., Trojet, S.N., Prere, M.-F., and Krisch, H.J.P.O. (2007). Phage-antibiotic synergy (PAS): β-lactam and quinolone antibiotics stimulate virulent phage growth. 2, e799.

Dedrick, R.M., Guerrero-Bustamante, C.A., Garlena, R.A., Russell, D.A., Ford, K., Harris, K., Gilmour, K.C., Soothill, J., Jacobs-Sera, D., and Schooley, R.T. (2019). Engineered bacteriophages for treatment of a patient with a disseminated drug-resistant Mycobacterium abscessus. Nature medicine 25, 730.

Diancourt, L., Passet, V., Verhoef, J., Grimont, P.A., and Brisse, S. (2005). Multilocus sequence typing of Klebsiella pneumoniae nosocomial isolates. J Clin Microbiol 43, 4178–4182.

Goren, M.G., Carmeli, Y., Schwaber, M.J., Chmelnitsky, I., Schechner, V., and Navon-Venezia, S. (2010). Transfer of carbapenem-resistant plasmid from Klebsiella pneumoniae ST258 to Escherichia coli in patient. Emerg Infect Dis 16, 1014–1017.

Gu, D., Dong, N., Zheng, Z., Lin, D., Huang, M., Wang, L., Chan, E.W.-C., Shu, L., Yu, J., and Zhang, R.J.T.L.I.D. (2018). A fatal outbreak of ST11 carbapenem-resistant hypervirulent Klebsiella pneumoniae in a Chinese hospital: a molecular epidemiological study. 18, 37–46.

Hardiman, C.A., Weingarten, R.A., Conlan, S., Khil, P., Dekker, J.P., Mathers, A.J., Sheppard, A.E., Segre, J.A., and Frank, K.M. (2016). Horizontal Transfer of Carbapenemase-Encoding Plasmids and Comparison with Hospital Epidemiology Data. Antimicrob Agents Chemother 60, 4910–4919.

Hung, C.H., Kuo, C.F., Wang, C.H., Wu, C.M., and Tsao, N. (2011). Experimental phage therapy in treating Klebsiella pneumoniae-mediated liver abscesses and bacteremia in mice. Antimicrob Agents Chemother 55, 1358–1365.

Koskella, B., and Meaden, S. (2013). Understanding bacteriophage specificity in natural microbial communities. Viruses 5, 806–823.

Kropinski, A.M., Mazzocco, A., Waddell, T.E., Lingohr, E., and Johnson, R.P. (2009). “Enumeration of Bacteriophages by Double Agar Overlay Plaque Assay,” in Bacteriophages: Methods and Protocols, Volume 1: Isolation, Characterization, and Interactions, eds. M.R.J. Clokie & A.M. Kropinski. (Totowa, NJ: Humana Press), 69–76.

Labrie, S.J., Samson, J.E., and Moineau, S. (2010). Bacteriophage resistance mechanisms. Nat Rev Microbiol 8, 317–327.

Letkiewicz, S., Międzybrodzki, R., Fortuna, W., Weber-Dąbrowska, B., and Górski, A.J.F.M. (2009). Eradication of Enterococcus faecalis by phage therapy in chronic bacterial prostatitis—case report. 54, 457.

Linkevicius, M., Sandegren, L., and Andersson, D.I. (2016). Potential of Tetracycline Resistance Proteins To Evolve Tigecycline Resistance. Antimicrob Agents Chemother 60, 789–796.

Mcvay, C.S., Velasquez, M., and Fralick, J.A. (2007). Phage therapy of Pseudomonas aeruginosa infection in a mouse burn wound model. Antimicrob Agents Chemother 51, 1934–1938.

Pendleton, J.N., Gorman, S.P., and Gilmore, B.F. (2013). Clinical relevance of the ESKAPE pathogens. Expert Rev Anti Infect Ther 11, 297–308.

Pitout, J.D., Nordmann, P., and Poirel, L. (2015). Carbapenemase-Producing Klebsiella pneumoniae, a Key Pathogen Set for Global Nosocomial Dominance. Antimicrob Agents Chemother 59, 5873–5884.

Pitout, J.D., Sanders, C.C., and Sanders Jr, W.E.J.T.a.J.O.M. (1997). Antimicrobial resistance with focus on β-lactam resistance in gram-negative bacilli. 103, 51–59.

Qi, Y., Wei, Z., Ji, S., Du, X., Shen, P., and Yu, Y.J.J.O.a.C. (2011). ST11, the dominant clone of KPC-producing Klebsiella pneumoniae in China. 66, 307–312.

Schooley, R.T., Biswas, B., Gill, J.J., Hernandez-Morales, A., Lancaster, J., Lessor, L., Barr, J.J., Reed, S.L., Rohwer, F., Benler, S.J.a.A., and Chemotherapy (2017). Development and use of personalized bacteriophage-based therapeutic cocktails to treat a patient with a disseminated resistant Acinetobacter baumannii infection. 61.

Seed, K.D., Faruque, S.M., Mekalanos, J.J., Calderwood, S.B., Qadri, F., and Camilli, A. (2012). Phase variable O antigen biosynthetic genes control expression of the major protective antigen and bacteriophage receptor in Vibrio cholerae O1. PLoS Pathog 8, e1002917.

Senchyna, F., Gaur, R.L., Sandlund, J., Truong, C., Tremintin, G., Kültz, D., Gomez, C.A., Tamburini, F.B., Andermann, T., Bhatt, A.J.D.M., and Disease, I. (2019). Diversity of resistance mechanisms in carbapenem-resistant Enterobacteriaceae at a health care system in Northern California, from 2013 to 2016. 93, 250–257.

Sulakvelidze, A., Alavidze, Z., and Morris, J.G., Jr. (2001). Bacteriophage therapy. Antimicrob Agents Chemother 45, 649–659.

Sumrall, E.T., Shen, Y., Keller, A.P., Rismondo, J., Pavlou, M., Eugster, M.R., Boulos, S., Disson, O., Thouvenot, P., and Kilcher, S.J.P.P. (2019). Phage resistance at the cost of virulence: Listeria monocytogenes serovar 4b requires galactosylated teichoic acids for InlB-mediated invasion. 15, e1008032.

Suttle, C.A. (2005). Viruses in the sea. Nature 437, 356–361.

Tan, D., Gram, L., Middelboe, M.J.A., and Microbiology, E. (2014). Vibriophages and their interactions with the fish pathogen Vibrio anguillarum. 80, 3128–3140.

Tétart, F., Repoila, F., Monod, C., and Krisch, H. (1996). “Bacteriophage T4 host range is expanded by duplications of a small domain of the tail fiber adhesin”. Elsevier).

Toth, A., Damjanova, I., Puskas, E., Jánvári, L., Farkas, M., Dobák, A., Böröcz, K., Paszti, J.J.E.J.O.C.M., and Diseases, I. (2010). Emergence of a colistin-resistant KPC-2-producing Klebsiella pneumoniae ST258 clone in Hungary. 29, 765–769.

Ventola, C.L. (2015). The antibiotic resistance crisis: part 1: causes and threats. P T 40, 277–283.

Yao, H., Qin, S., Chen, S., Shen, J., and Du, X.-D.J.T.L.I.D. (2018). Emergence of carbapenem-resistant hypervirulent Klebsiella pneumoniae. 18, 25.

